# Environmental enrichment alleviates the deleterious effects of stress in experimental autoimmune encephalomyelitis

**DOI:** 10.1101/2020.04.10.033662

**Authors:** Antoine Philippe Fournier, Erwan Baudron, Isabelle Wagnon, Philippe Aubert, Denis Vivien, Michel Neunlist, Isabelle Bardou, Fabian Docagne

## Abstract

Clinical observations support the hypothesis that stressful events increase relapse occurrence in multiple sclerosis patients, while stress-reduction strategies can modulate this effect. However, a direct cause-effect relationship between stress level and relapse cannot be firmly established from these data. The purpose of this work was to address whether modulation of stress could interfere with symptom relapse in an animal model of multiple sclerosis with relapsing-remitting course. We report that repeated acute stress induced a twofold increase in relapse incidence in experimental autoimmune encephalomyelitis. On the other hand, environmental enrichment reduced relapse incidence and severity, and reversed the effects of repeated acute stress. These data provide the platform for further studies on the biological processes that link stress and multiple sclerosis relapses in a suitable animal model.

## INTRODUCTION

Multiple sclerosis (MS) is a chronic inflammatory disease of the central nervous system that leads to various neurological manifestations, including sensitive and motor symptoms. In the most frequent form, the relapsing-remitting subtype, these symptoms occur in an unpredictable manner during periods of relapses, followed by periods of remissions when signs of disease activity are reduced or absent. There is considerable interest in identifying the factors that lead to the occurrence of relapse.

Clinical observations have evidenced a link between acute stress and occurrence of relapses in MS patients. Several studies report that acute stressful life events (marital separation, difficulties at work, changes in place of residence, death of a relative, etc.)^1^ increase the probability for MS patients to experience a relapse^2^. A prospective study showed that three or more stressful life events in a four week period led to a fivefold increase of MS relapse rate, and that the presence of at least one stressful life events is sufficient to increase by threefold the rate of relapse during the following four weeks^3^. Conversely, stress reduction strategies have shown positive effects in MS patients^4^.

Considering these clinical observations, several studies have attempted to reproduce the effects of stress in animal models of MS. In most cases, stress was shown to reduce, rather than increase, the severity of experimental autoimmune encephalomyelitis (EAE)^5^. However, most of the studies have used chronic stress (more than one hour per day for more than five days)^5^, which has long been known to inhibit immune response, rather than acute stress events. One study reported an advance in the onset of EAE after repeated acute stress^6^. More recently, mild chronic stress was shown to exacerbate and accelerate the clinical symptoms of rat EAE^7^.

However, in all of these earlier studies, the stress paradigm was applied in the early phase, prior to symptom onset, and not during the remission phase. Thus, these studies did not allow addressing the effect of stress on relapses in relapsing-remitting context.

Overall, the question remains of whether acute stress, when occurring during remission periods, can influence the incidence and severity of relapses. In addition, the effect of stress reduction on relapses was never tested in relapsing-remitting animal models of MS. To answer these questions, we set up a paradigm of repeated acute stress applied during the remission phase of relapsing-remitting EAE. We observed that stress exposure precipitated relapse, so that in stressed animals, the incidence of relapse was doubled, and the delay before the occurrence of relapse was significantly reduced. Conversely, environmental enrichment, previously shown to reduce stress in rodents, reduced relapse incidence and severity, and reversed the effects of stress.

## METHODS

### Animals

Experiments were performed on female SJL/J mice (Janvier, Le Genest-Saint-Isle, France) maintained under specific pathogen-free conditions at the Centre Universitaire de Ressources Biologiques (Basse-Normandie, France). This study and the procedures thereof were approved by the French ministry of education and research (Project licence APAFIS#2887-2015112017418114v2; Center agreement #D14118001) in accordance with the French (Decree 87/848) and European (Directive 86/609) guidelines. Animals were monitored once a day for signs of pain, posture, reactivity, activity, signs of distress, food/drink consumption and follow-up of body weight. Humane euthanasia was planned to be applied to any animal showing at least one of the following signs: prostration, body weight loss > 20%, lack of reaction, prolonged inactivity or other signs of distress, stop of food and/or drink consumption.

### Experimental autoimmune encephalomyelitis (EAE)

*Relapsing remitting EAE* (*PLP-induced EAE*) was induced in 8-week-old female SJL/J mice via subcutaneous immunization with 200 μg recombinant myelin proteolipid protein (PLP_139-151_, Eurogentec) in an emulsion mix (volume ratio 1:1) with Complete Freund’s Adjuvant (CFA; Difco Laboratories) containing 800 μg of heat-killed Mycobacterium tuberculosis H37Ra (MBT; Difco). The emulsion was administered to regions above the shoulders and the flanks (total of 4 sites; 50 μl at each injection site). All animals were injected intraperitoneally with 200 ng pertussis toxin derived from Bordetella pertussis (Sigma-Aldrich) in 200 μL saline at the time of, and after 48 hours following immunization. EAE induction was performed with the application of local analgesia (lidocaine patch) to prevent discomfort and pain. The animals were euthanized 80 days after EAE induction under deep anesthesia (5% isoflurane, O2/N2O 1/1) by exsanguination via intracardiac infusion of saline.

*Clinical score:* mice were examined daily for clinical signs of EAE and were scored as followed: 0, no disease; 1, limp tail; 2, hindlimb weakness; 3, complete hindlimb paralysis; 4, hindlimb paralysis plus forelimb paralysis; and 5, moribund or dead. All clinical score were assessed daily by an examiner blinded to the EAE group. A relapse was defined as a sustained increase (minimum duration of 2 days) of at least 0.5 in clinical score.

### Water avoidance stress (WAS)

WAS (one hour per day during four days) was performed by placing asymptomatic EAE mice on a platform (4cm diameter) positioned at the center of a plastic container filled with water at room temperature up to 1 cm below the top of the platform. EAE mice were randomly assigned to the two experimental groups. Non-WAS mice were handled identically, but placed in an empty and clean cage.

### Environmental enrichment (EE)

Enriched mice were housed in Marlau^™^ cages [Viewpoint; 58 × 40 × 32 cm (L × W × H)], which allowed social interactions (15 mice per cage). These cages are composed of two floors. The ground floor has two compartments: one containing food, the other one containing water bottles, three running wheels and an elevated nest. The upper floor contains a maze changed three times a week. Because of one-way doors between the two ground compartments, mice had to climb to the upper floor using a ladder, pass through the maze and go down to the food compartment using a slide tunnel to reach food. Enriched mice were housed in these Marlau™ cages 14 days prior to EAE induction until the end of the experiment. Non-enriched animals were placed in standard cages (STD) with ad libitum access to food and water.

### Statistical analysis

Results are presented as the mean ± SEM. Normality tests were performed on all samples (D’Agostino-Pearson omnibus test and Shapiro-Wilk test). When normality could not be assumed, we used non-parametric tests (Mann-Whitney’s U-test), which are the most stringent in these conditions. For the comparison of the number of feces in STD, STD+WAS, EE and EE+WAS groups, we used a Kruskall-Wallis test followed by a Dunn’s multiple comparison as a *post hoc* test. Onset and relapse incidence curves were analysed with Gehan-Breslow-Wilcoxon test. Data were analysed using GraphPad Prism 7.0. Two groups were considered to be significantly different when *P*< 0.05.

## RESULTS

Repeated acute stress has been defined as a short exposure to stressing stimulus (<1h) repeated once daily during less than 5 days (as opposed to repeated chronic stress: >1h during more than 5 days)^5^. Here, to induce repeated acute stress, we applied a paradigm in which mice were exposed to water avoidance stress (WAS) once a day during 4 days during the remission period of EAE (Figure 1A-B). The stressful nature of this paradigm was assessed by measuring fecal pellet output during the exposure to stress (Figure 1C), as a measure of stress-induced colonic motility^8^. This procedure is fully non-invasive and was preferred to other methods such as measurement of stress hormones in the blood-stream that require invasive procedures and thus constitute a source of stress for the animals.

**Figure 1:**
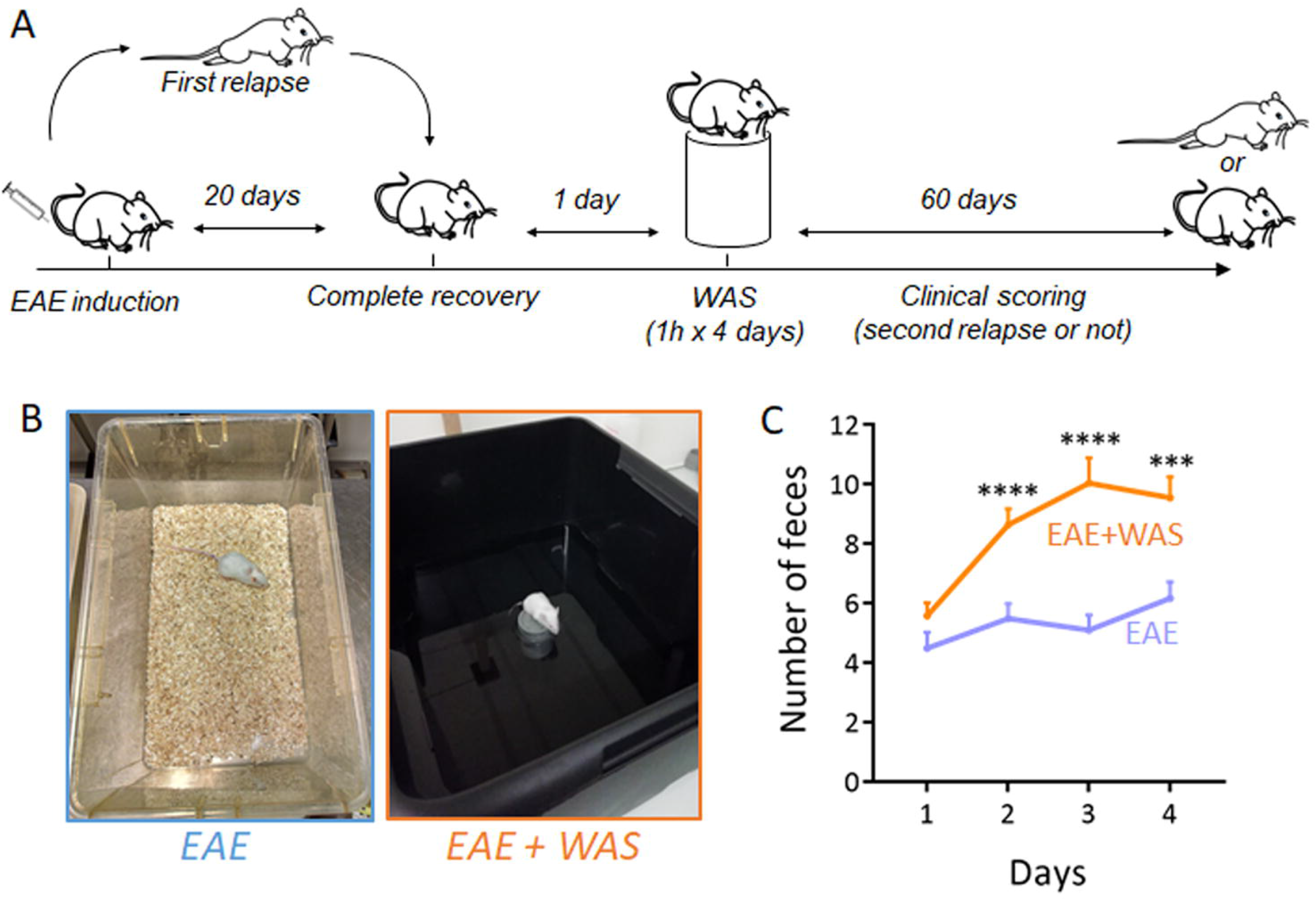
Experimental design for water avoidance stress. **(A)** Water avoidance stress (WAS, one hour per day for four days) was applied during the remission phase of PLP-induced relapsing-remitting EAE mice and the clinical scoring was assessed during 60 days. **(B)** Photography of the experimental system. Mice were placed for 1 hour on a platform positioned at the center of a plastic container filled with water at room temperature up to 1 cm below the platform level. Mice were randomly assigned to the two experimental groups. Non-WAS mice were handled identically, but placed in a standard cage. **(C)** Measure of fecal pellet outputs in the two experimental groups as an index of stress-induced colonic motility (n=26 and n= 28 for EAE and EAE+WAS groups respectively). \*\*\**P*<0.001; \*\*\*\**P*<0.0001, Mann Whitney’s U-test.

The exposure to stress induced an increase in clinical score which reached significance at days 43, 44 and 46, and between days 58 and 63 (Figure 2A). These differences in mean clinical score along the course of EAE were the result of two parameters. First, relapses occurred more often in stressed animals, so that the incidence of relapses was increased by 1.58-fold at day 75 (Figure 2B). Second, relapses occurred earlier in stressed animals (mean day of relapse onset: 40.53±3.43 vs. 55.33±4.92; *P*=0.0189; Table 1). Noteworthy, the severity of EAE was not affected by stress (mean peak score: 2.77 ± 0.29 vs. 2.50 ± 0.47. Table 1).

**Figure 2:**
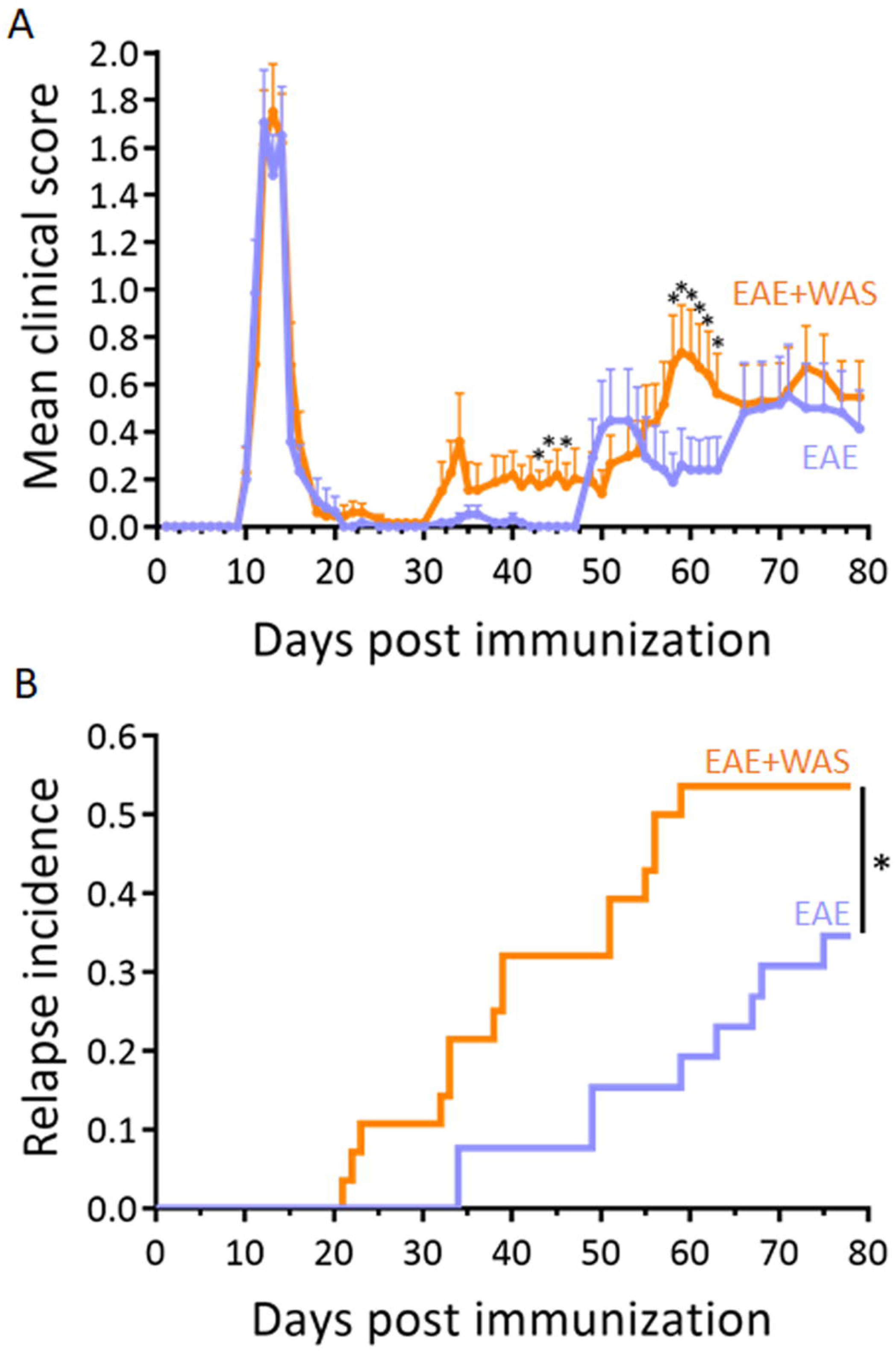
Acute stress precipitates relapse in EAE animals. **(A)** Clinical score was assessed daily by an examiner blinded to the treatment (n=26 and n= 28 for EAE and EAE+WAS groups respectively; \**P*<0.05). **(B)** Clinical evaluation expressed as the long-term relapse incidence in EAE and EAE+WAS groups (\**P*<0.05, Gehan-Breslow-Wilcoxon test).

**Table 1:**
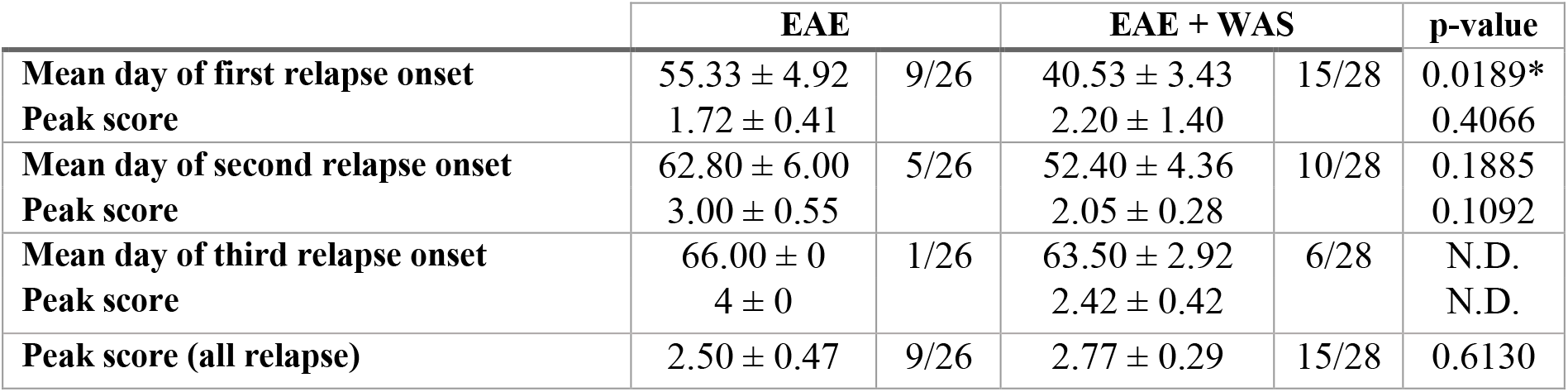
Values for mean day of relapse onset and mean peak clinical score in the EAE and EAE+WAS groups. P values are given for t-test. \**P*<0.05.

|  | EAE |  | EAE + WAS |  | p-value |
| --- | --- | --- | --- | --- | --- |
| Mean day of first relapse onset | 55.33 ± 4.92 | 9/26 | 40.53 ± 3.43 | 15/28 | 0.0189* |
| Peak score | 1.72 ± 0.41 | | 2.20 ± 1.40 | | 0.4066 |
| Mean day of second relapse onset | 62.80 ± 6.00 | 5/26 | 52.40 ± 4.36 | 10/28 | 0.1885 |
| Peak score | 3.00 ± 0.55 | | 2.05 ± 0.28 | | 0.1092 |
| Mean day of third relapse onset | 66.00 ± 0 | 1/26 | 63.50 ± 2.92 | 6/28 | N.D. |
| Peak score | 4 ± 0 | | 2.42 ± 0.42 | | N.D. |
| Peak score (all relapse) | 2.50 ± 0.47 | 9/26 | 2.77 ± 0.29 | 15/28 | 0.6130 |

Environmental enrichment has been described as an efficient paradigm to reduce stress in laboratory rodents ^9^. Here, we bred mice in standard or enriched environment prior to EAE induction and exposure to repeated acute stress (Figure 3A-B). This paradigm of standard enrichment reversed the increase of fecal pellet output (used as index of stress-induced colonic motility) induced by WAS (Figure 3C).

**Figure 3:**
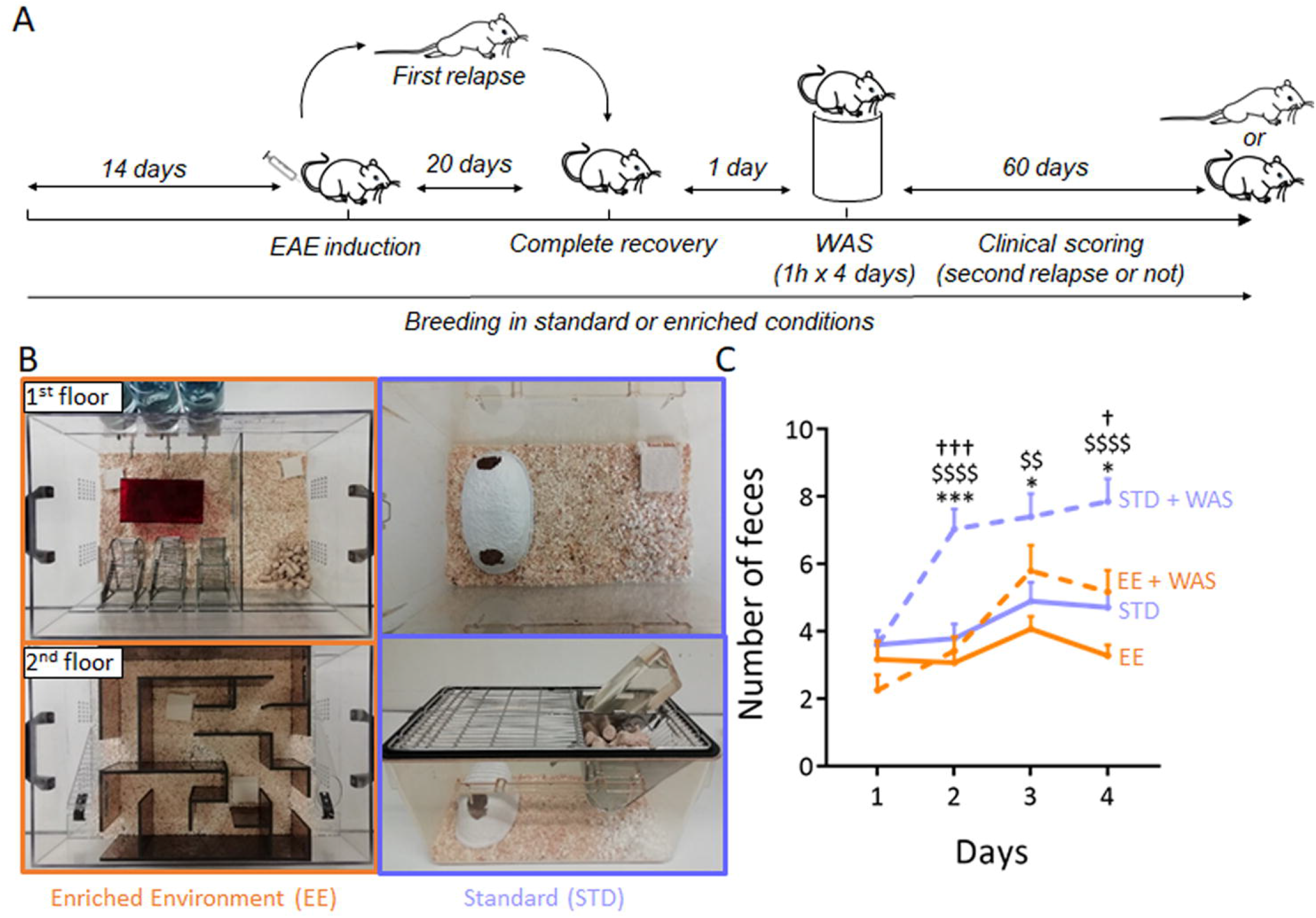
Experimental design for environment enrichment. **(A)** Environmental enrichment (EE) was applied from 14 days pre-EAE induction to the end of the protocol. Standard (STD) animals were kept in standard breeding conditions. WAS was applied to half of the animals of each group during the remission phase of EAE. In total, 4 experimental groups were studied: STD, STD+WAS, EE, EE+WAS. (B) Photography of the STD and EE breeding cages (see methods for details). **(C)** Measure of fecal pellet outputs in the four experimental groups as an index of stress-induced colonic motility (n=16, n= 21, n=12 and, n=12 for STD, STD+WAS, EE, and EE+WAS groups respectively; \**P*<0.05; \*\*\**P*<0.001, STD vs. STD+WAS; ^$$$$^*P*<0.0001, EE vs. STD+WAS; ^†††^*P*<0.001, EE+WAS vs. STD+WAS; Kruskall-Wallis test + Dunn’s multiple comparison).

Strikingly, animals bred in enriched environment showed a reduction in the first symptomatic peak of EAE (Days 10-14 post immunization, Figure 4A) and a drastic reduction of clinical score during the relapse phase (days 49-80, Figure 4A). These differences in mean clinical score along the course of EAE were the result of several parameters: First, the incidence of the disease and the incidence of relapse were reduced in enriched animals (Figure 4B-C), so that the incidence of disease was decreased by 1.52-fold at day 25 (Figure 4B) and the incidence of relapse was decreased by 2.06-fold at day 70 (Figure 4C). Second, the onset of the disease tended to occur later in enriched animals (mean day of relapse onset: 14.67±0.41 vs. 13.74±0.35 vs.; *P*=0.0581; Table 2).

**Figure 4:**
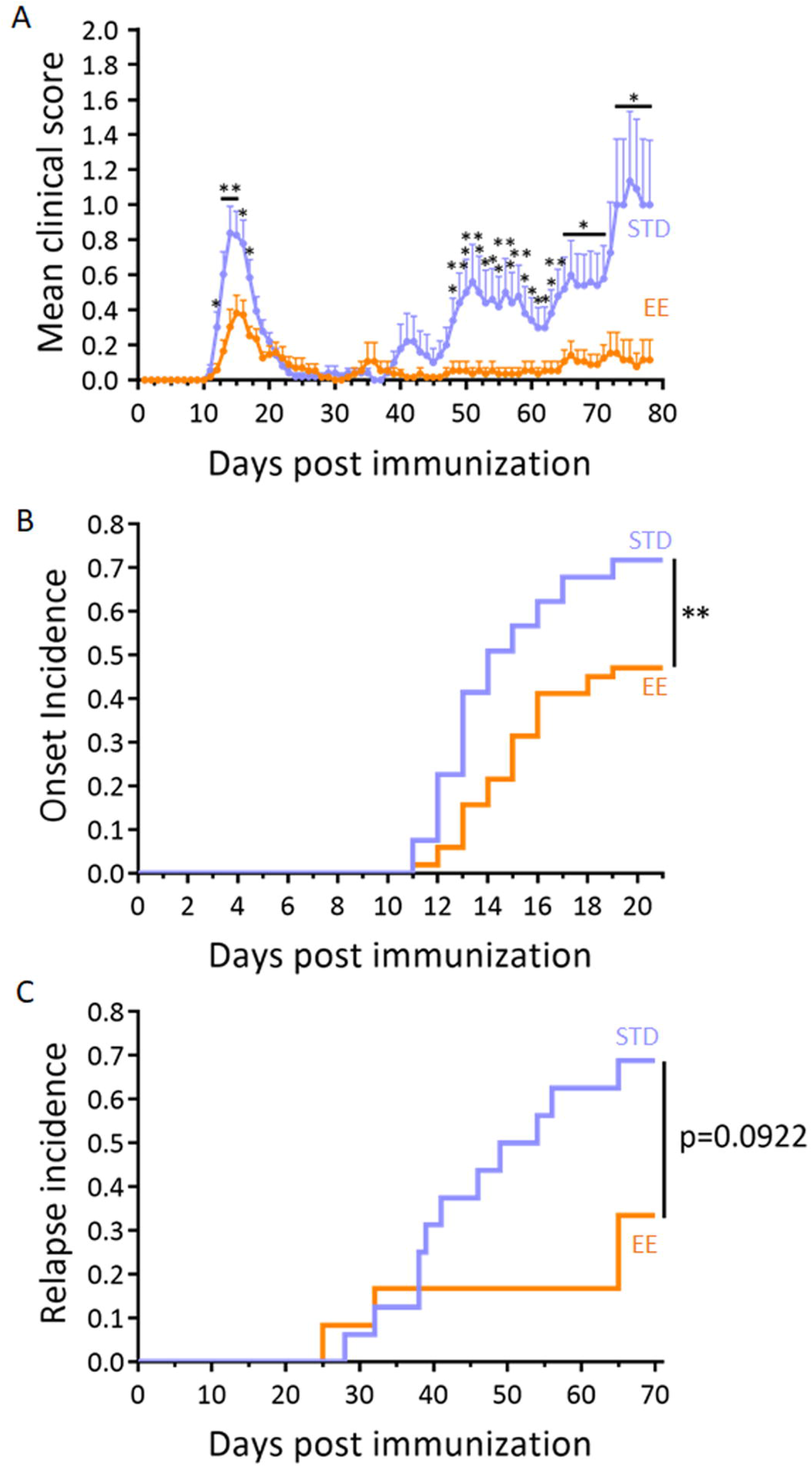
Environmental enrichment reduced EAE severity. **(A)** Clinical score assessed daily by an examiner blinded to the treatment in the standard (STD) and enriched environment (EE) groups (n=53 and n=51 for STD and EE groups respectively; \**P*<0.05; \*\**P*<0.01, Mann-Whitney’s U-test). **(B)** Clinical evaluation expressed as the EAE onset incidence in EE and STD groups (\*\**P*<0.01, Gehan-Breslow-Wilcoxon test). **(C)** Clinical evaluation expressed as the long-term EAE relapse incidence in EE and STD groups (n=16 and n=12 for STD and EE groups respectively; *P*=0.0922, Gehan-Breslow-Wilcoxon test).

**Table 2:** Values for mean day of onset, mean day of relapse and mean peak clinical score in the STD or EE groups. P values are given for t-test or, when normality could not be assumed, Mann-Whitney’s U-test.

|  | STD |  | EE |  | p-value |
| --- | --- | --- | --- | --- | --- |
| Mean day of onset | 13.74 ± 0.35 | 38/53 | 14.67 ± 0.41 | 24/51 | 0.0581 |
| Peak score | 1.64 ± 0.17 | | 1.27 ± 0.16 | | 0.2070 |
| Mean day of first relapse onset | 44.18 ± 3.33 | 11/16 | 46.75 ± 10.63 | 4/12 | 0.7597 |
| Peak score | 1.64 ± 0.44 | | 1.63 ± 0.47 | | 0.5963 |
| Mean day of second relapse onset | 55.29 ± 4.11 | 7/16 | 47.00 ± 0 | 1/12 | N.D. |
| Peak score | 2.43 ± 0.46 | | 1.50 ± 0 | | N.D. |
| Mean day of third relapse onset | 65.00 ± 2.42 | 4/16 | 69.00 ± 0 | 1/12 | N.D. |
| Peak score | 2.63 ± 0.55 | | 1.50 ± 0 | | N.D. |
| Peak score (all relapse) | 2.77 ± 0.37 | 11/16 | 1.63 ± 0.47 | 4/12 | 0.1143 |

Considering that repeated acute stress increased relapse rate and that environmental enrichment reduced the severity of EAE, our next question was to ask as to whether environmental enrichment may reduce the effects of stress. Mice bred in standard or enriched environment were subjected to WAS during the remission period (Figure 3A). In agreement with above results, mice bred in enriched environment showed a reduction in clinical score during the first peak of disease (day 11 to 15, Figure 5A). The mean clinical score gradually increased after the remission period in standard animals, but not in enriched animals (Figure 5A), reaching statistical difference from day 43 post immunization to the end of the experiment (day 80). This difference was the reflect of a dramatic difference in the incidence of relapse (71.4% vs. 8.3%, *P*=0.0011; Figure 5B). Only one animal (from n=12) suffered a relapse in the enriched group, and that relapse occurred much later than what observed in the standard group (61 days vs. a mean of 43.73±3.15 in standard animals; Table 3). The peak score of this relapse was also reduced as compared to the mean of the standard group (1.5±0 vs. 2.00±0.32; Table 3). In addition, although 60% of the animals with a first relapse (9 animals out of 15) also suffered a second relapse later in the standard group (58.43±4.28 days post immunization; Table 3), no second relapse was observed in the enriched group.

**Figure 5:**
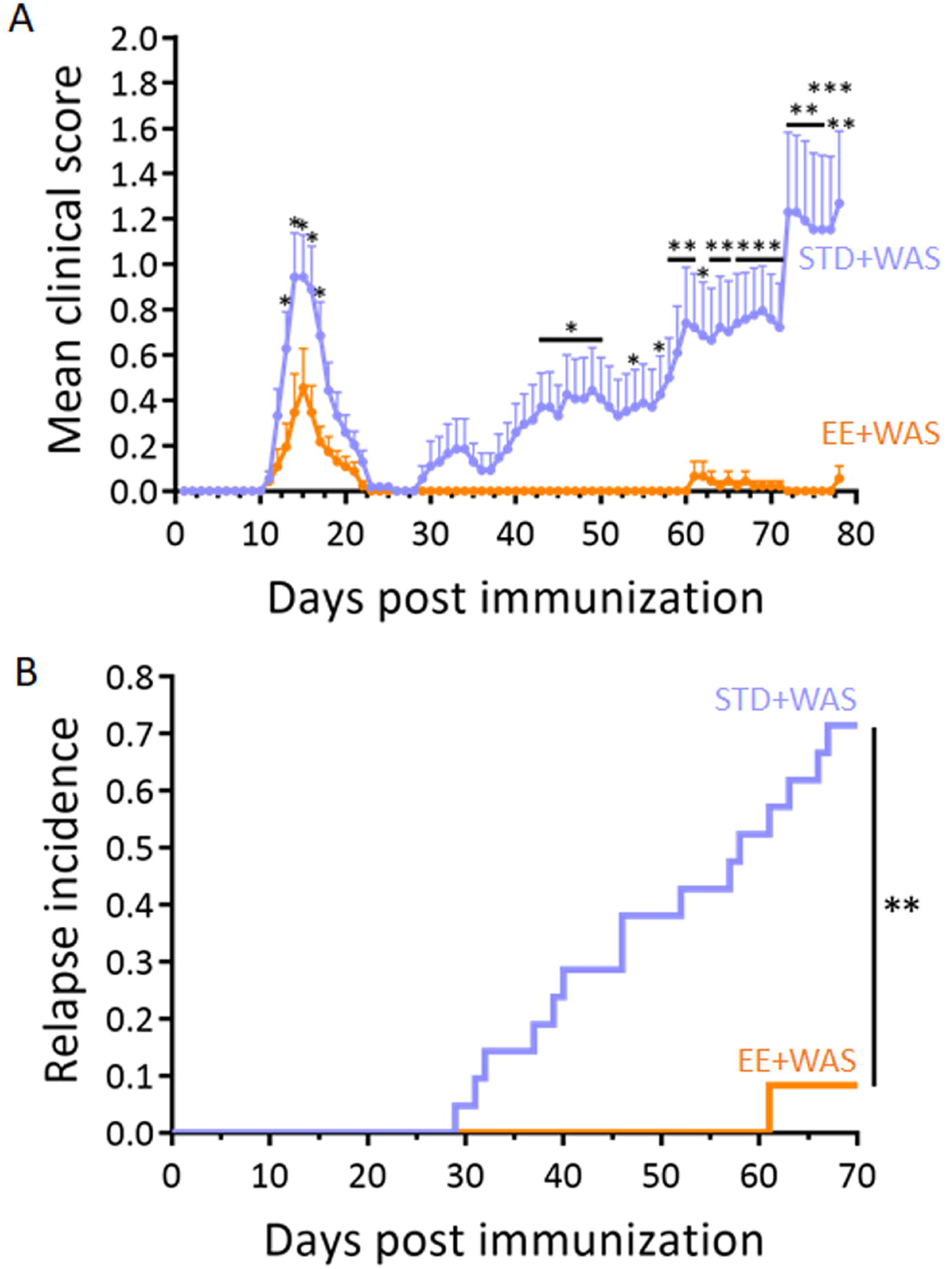
Environmental enrichment reversed the effects of stress on EAE severity. **(A)** Clinical score assessed daily by an examiner blinded to the treatment in the STD+WAS and EE+WAS groups (n=21 and n=12 for STD+WAS and EE+WAS groups respectively; \**P*<0.05; \*\**P*<0.01; \*\*\**P*<0.001, Mann Whitney’s U-test). **(B)** Clinical evaluation expressed as the longterm EAE relapse incidence in STD+WAS and EE+WAS groups (n=21 and n=12 for STD+WAS and EE+WAS groups respectively; **P<0.01, Gehan-Breslow-Wilcoxon test).

**Table 3:** Values for mean day of relapse onset and mean peak clinical score in the STD+WAS and EE+WAS groups.

|  | STD+WAS |  | EE+WAS |  | p-value |
| --- | --- | --- | --- | --- | --- |
| Mean day of first relapse onset | 43.73 ± 3.15 | 15/21 | 61.00 ± 0 | 1/12 | N.D. |
| Peak score | 2.00 ± 0.32 | | 1.50 ± 0 | | N.D. |
| Mean day of second relapse onset | 58.43 ± 4.28 | 9/21 | N.D. | 0/12 | N.D. |
| Peak score | 2.36 ± 0.31 | | N.D. | | N.D. |
| Mean day of third relapse onset | 56.00 ± 2.00 | 2/21 | N.D. | 0/12 | N.D. |
| Peak score | 2.50 ± 1.00 | | N.D. | | N.D. |
| Mean day of fourth relapse onset | 76.00 ± 0 | 1/21 | N.D. | 0/12 | N.D. |
| Peak score | 1.00 ± 0 | | N.D. | | N.D. |
| Peak score (all relapse) | 2.50 ± 0.30 | 15/21 | 1.50 ± 0 | 1/12 | N.D. |

**Abbreviations:** MS: Multiple sclerosis; EAE: Experimental autoimmune encephalomyelitis; RR-EAE : Relapsing-Remitting EAE ; PLP : Proteolipid protein ; CFA : Complete Freund adjuvant ; WAS : Water avoidance stress; EE : Enriched Environment ; STD : Standard Environment

## DISCUSSION

The present study overall shows that repeated acute stress precipitates relapses in RR-EAE and that environmental enrichment, previously reported to reduce stress in laboratory rodents^9^, prevents the occurrence of relapses and alleviates the effect of repeated acute stress on RR-EAE severity.

Controversial results have been reported in the past concerning the effects of stress on EAE^10^. A general consensus has emerged to state that chronic stress ameliorates, while acute stress worsens, disease course. The outcome of these studies was generally the evolution of clinical score in a chronic, monophasic model, rather than the effect on relapses in a relapsing-remitting model, as described here. The data presented here show that repeated acute stress during the remission phase of RR-EAE is sufficient to precipitate relapses in absence of other inducers. These data complement previous reports of an exacerbation of disease by chronic mild stress^7^. Our data match clinical observation in MS patients, previously summarized in a meta-analysis^11^ and describing that stressful life events exacerbate MS. The paradigm developed in the present study may therefore be used to investigate the mechanisms responsible for stress-induced occurrence of relapses.

The two major stress response systems, the hypothalamic-adrenal-pituitary axis and the sympathetic nervous system, have been suggested to mediate the effects of stress on relapses^12^. In addition to these pathways, stress also increases intestinal barrier permeability^13^. Animal studies give a possible link between gut inflammation and MS: intestinal barrier dysfunction occurs at the onset of monophasic EAE^14^ and a pro-inflammatory response in the gut has been shown to trigger EAE^15,16^. Conversely, the sequestration of Th17 cells in the gut confers resistance to EAE^17^. Together, these studies suggest a sequence of events involving the gut in the pathogenesis of EAE: (i) initial expansion and activation of T-cell in gut-associated lymphoid tissues, leading to (ii) recruitment of auto-antibody-producing B cells and (iii) migration to brain draining cervical nodes to finally trigger autoimmune encephalomyelitis. Increased permeability of the intestinal barrier is a key feature of gut inflammation^18^ and could thus initiate this cascade of events, which may potentiated by stress. In addition, stress may also induce neurovascular pathology, leading to BBB permeation^19,20^ which could provide an additional explanation for the effect of stress in EAE. The paradigm described in the present work could be useful to test these mechanistic hypotheses in further studies.

While the effects of stress on RR-EAE disease described here corroborate previous reports in experimental models^7^ and coincide with clinical data^11^, more intriguing and novel are our data concerning the effects of environmental enrichment. Indeed, we show that animals bred in enriched environment show a reduced severity and incidence of the first RR-EAE peak, and a drastic amelioration of the second phase of the disease. In addition, environmental enrichment alleviated the effects of repeated acute stress, applied during the remission phase, on the incidence of relapse. These data suggest that animals bred in an enriched environment have lost their sensitivity to repeated acute stress on RR-EAE symptoms. The fact that environmental enrichment reduces the severity of RR-EAE from the first peak suggests that this breeding regimen exert a “conditioning” effect on animals towards a lesser sensitivity to EAE. Interestingly, although repeated acute stress prompts standard animals towards a higher rate of relapse, this stress paradigm does not compensate for the beneficial “conditioning” effect brought by environmental enrichment.

The mechanism by which environmental enrichment alleviates the effects of stress is also an intriguing question. Previous report suggested that environmental enrichment increases cerebral activity in the prefrontal cortex and that this increased activity participates in setting up stress resiliency^21^. This suggests that environmental enrichment could drive a manner of cerebral training in rodents that would lead to the reported beneficial effects. However, one cannot exclude that this regimen may reverse the effects of stress by other-though not necessarily exclusive-ways. Several positive effects of environmental enrichment, in addition to cerebral training, may account for its beneficial effects, including increase in physical activity, or changes in metabolic parameters^22^.

In conclusion, this study shows that repeated acute stress increases relapse rate and incidence in an experimental paradigm relevant to relapsing-remitting MS, the most frequent form of this disease. These data in animal models have to be seen in the light of recent successful attempts to use stress management strategies as an adjunct to the available MS treatments^23,24^. The use of preclinical animal models may provide rationales and mechanistic explanations for the use of these strategies in MS management.

## Abbreviations

MS: Multiple sclerosis
EAE: Experimental autoimmune encephalomyelitis
RR-EAE: Relapsing-Remitting EAE
PLP: Proteolipid protein
CFA: Complete Freund adjuvant
WAS: Water avoidance stress
EE: Enriched Environment
STD: Standard Environment

## Ethics approval

This study and the procedures thereof were approved by the French ministry of education and research (Project reference APAFIS#2887-2015112017418114v2; Center agreement #D14118001) in accordance with the French (Decree 87/848) and European (Directive 86/609) guidelines.

## Availability of data and materials

The datasets used and analysed during the current study are available from the corresponding author.

